# Polinton-like viruses and virophages are widespread in aquatic ecosystems

**DOI:** 10.1101/2019.12.13.875310

**Authors:** Christopher M. Bellas, Ruben Sommaruga

## Abstract

Polintons are virus-like transposable elements found in the genomes of eukaryotes that are considered the ancient ancestors of most eukaryotic dsDNA viruses^1,2^. Recently, a number of Polinton-Like Viruses (PLVs) have been discovered in algal genomes and environmental metagenomes^3^, which share characteristics and core genes with both Polintons and virophages (Lavidaviridae)^4^. These viruses could be the first members of a major class of ancient eukaryotic viruses, however, only a few complete genomes are known and it is unclear whether most are free viruses or are integrated algal elements^3^. Here we show that PLVs form an expansive network of globally distributed viruses, associated with a range of eukaryotic hosts. We identified PLVs as amongst the most abundant individual viruses present in a freshwater lake virus metagenome (virome), showing they are hundreds of times more abundant in the virus size fraction than in the microbial one. Using the major capsid protein genes as bait, we retrieved hundreds of related viruses from publicly available datasets. A network-based analysis of 976 new PLV and virophage genomes combined with 64 previously known genomes revealed that they represent at least 61 distinct viral clusters, with some PLV members associated with fungi, oomycetes and algae. Our data reveals that PLVs are widespread in predominantly freshwater environments and together with virophages, represent a broad group of eukaryotic viruses which share a number of genes.

## Main text

Virophages infect microbial eukaryotes, but are entirely dependent on co-infecting giant viruses to carry out their replication^5^. Some are parasitic, decreasing giant virus replication and increasing host cell population survival rates^6,7^. One such virus is Mavirus^8^, which integrates into the genome of the marine flagellate *Cafeteria roenbergensis*, where it acts as an antivirus defense system against the giant virus CroV^7^. All virophages possess relatively small genomes of 17-30 kb, with a core set of genes that include a Major Capsid Protein (MCP), Minor Capsid Protein (mCP) and DNA packaging ATPase^5^. However, Mavirus is also clearly related to the Polintons^8^, both in ability to integrate into a eukaryotic host and in possession of a protein-primed type B DNA polymerase (pPolB) and a retroviral-like (RVE family) integrase gene, which together are core Polinton genes. Hence a Mavirus-like virus could be the ancestral virophage^1^, however such conclusions are derived from only two known Mavirus-like genomes^8,9^. The recent discovery of ca. 25 Polinton-Like Virus (PLV) genomes^3,10,11^ has revealed a further group of related and presumed algal viruses, with several being associated with, or even integrated into, giant virus genomes which infect algae^10,11^. Only one PLV, *Tetraselmis striata* virus (originally named *Tetraselmis viridis* virus)^12^ has been observed as a virus particle, the remainder were discovered in sequencing projects or database searches based on the novel capsid gene of an unusual giant virus associated element, *Phaeocystis globosa* virus virophage (PgVV)^3^. When further capsid genes are identified, it therefore probable that many more PLVs will be discovered in sequencing datasets.

Throughout, we refer to the above entities which are similar in length and genomic content, all possessing a double jelly-roll fold MCP, an mCP and ATPase gene. Hence we define a Polinton as an integrated, probably viral, element in a eukaryotic genome that also encodes an RVE family integrase and pPolB gene; a Polinton-like virus (PLV) as a virus of similar size and genomic structure, found in metagenomes or integrated into a genome, which contains the three core genes above plus a varied compliment of additional genes (but without co-occurring pPolB and RVE); a virophage as any entity described above that possess a virophage type MCP gene (BLASTP E-value <10^-5^ to known virophages). Despite this categorisation, there is still considerable overlap between the groups and by this definition Mavirus is both a Polinton and virophage.

In a search for giant viruses in oligotrophic, alpine lakes^13^, we created paired virus-size fraction (< 0.2 μm) and microbial metagenomes (> 0.2 μm) from the high mountain lake, Gossenköllesee (2417 m above sea level, Austria) generating 127 GB of metagenomic sequencing data, which we assembled into contigs (see Methods). During annotation of the dataset, we discovered 32 new virophage genomes, small contigs (10-40 kb) with virophage type MCP genes (Table S1). Additionally, we identified many relatively abundant virus contigs with small circular genomes, or containing terminal repeats, that shared several homologous genes with virophages, but possessed no detectable virophage MCP gene (see Methods). We reasoned these contigs could be PLVs considering the genomic similarities to those identified previously^3^ (Table 1). To confirm this, we annotated the most abundant suspected PLV contig using remote protein homology detection via HHpred^14^ against the Protein Data Bank^15^ to identify an ATPase, minor capsid protein (mCP) and an MCP possessing a double jelly-roll fold and distant homology to the giant virus *Paramecium bursaria Chlorella* virus (PBCV-1) MCP gene (Table S2). This confirmed it as a PLV, as a similar double jelly-roll fold MCP gene has previously been identified in Polintons and PLVs^3,16^. We retrieved all related viruses from our metagenomes using this MCP gene as a query (BLASTP Evalue 10^-5^) before annotating the next suspected PLV and repeating the process. Each time a new MCP gene was identified, it was used to retrieve similar MCPs from the metagenome until all suspected PLVs had an annotated MCP gene (Table S1) or were fully annotated by HHpred (Table S2). To complete our search, we then retrieved all contigs that possessed a gene homologous to any previously described Polinton or PLV MCP gene from the literature^3,16^ (BLASTP E-value cutoff 10^-5^).

**Table 1.**
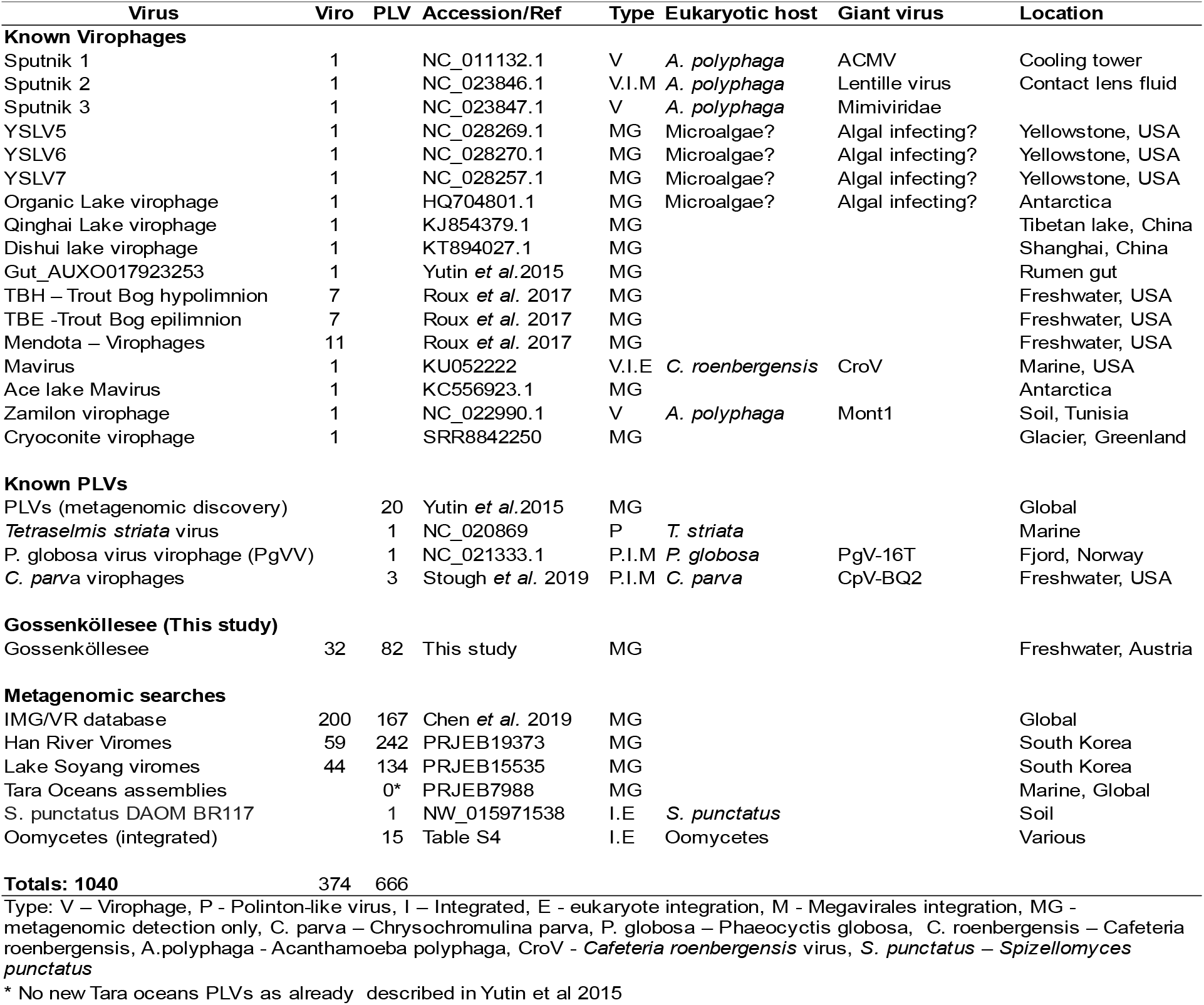
PLV and virophage contigs used and detected in this study (10-42 kb).

Despite identification of ATPase and mCP genes via HHpred, many suspected PLV contigs remained without an identifiable MCP gene (Table S2). In such cases, we carried these contigs forward in the analysis and identified the MCP at a later stage via alignment of core proteins clusters, which were used as a more sensitive HHpred query (Methods). Our putative PLV dataset was then subject to confirmation via analysis of shared gene content with known PLVs and virophages (Methods). This confirmed 82 new PLVs from Gossenköllesee, many of which appeared to represent complete genomes based on the presence of terminal repeats or a circular contig (Table S1). We also retrieved 16 related elements which we detected in eukaryotic genomes (Table S3) via a BLASTP search of the Gossenköllesee MCP genes against the GenBank non-redundant protein database. One element was found in the soil fungus *Spizellomyces punctatus* and 15 putative complete genomes were retrieved from oomycetes. In the fungal genome and most oomycetes, these elements were integrated into the middle of large contigs of the eukaryotic genome.

To determine the relationship between the new PLVs found in Gossenköllesee and those already described, we constructed a maximum likelihood tree of the MCP genes, including MCP identified in known Polintons^16^ (n=56) and PLV (n=25) (Table 1) along with more distantly related homologs from GenBank, which we identified via an iterative PSI-BLAST search based on our new capsid genes (Methods). The majority of Gossenköllesee MCPs formed novel groups, several of which had little homology to previously described Polinton or PLV capsid homologs (Figure 1). These included the GKS-1 group, which was composed of seven Gossenköllesee PLVs and homologues MCPs detected in oomycetes genomes; the GKS 2 group, which was composed of 16 Gossenköllesee MCP genes and had no known integrated representatives; and the VC20 group, a novel group of five viruses from Gossenköllesee (Virus cluster 20; Figure 1) which has identifiable MCP homologs in amoeba. An alignment of the MCP genes from the VC20 group used as a HHpred query detected distant homology to Singapore grouper iridovirus MCP (90.2% confidence), hence members of this VC20 group were distinct from known Polintons, PLVs or indeed virophages, which are also known to co-infect amoebae.

**Figure 1.**
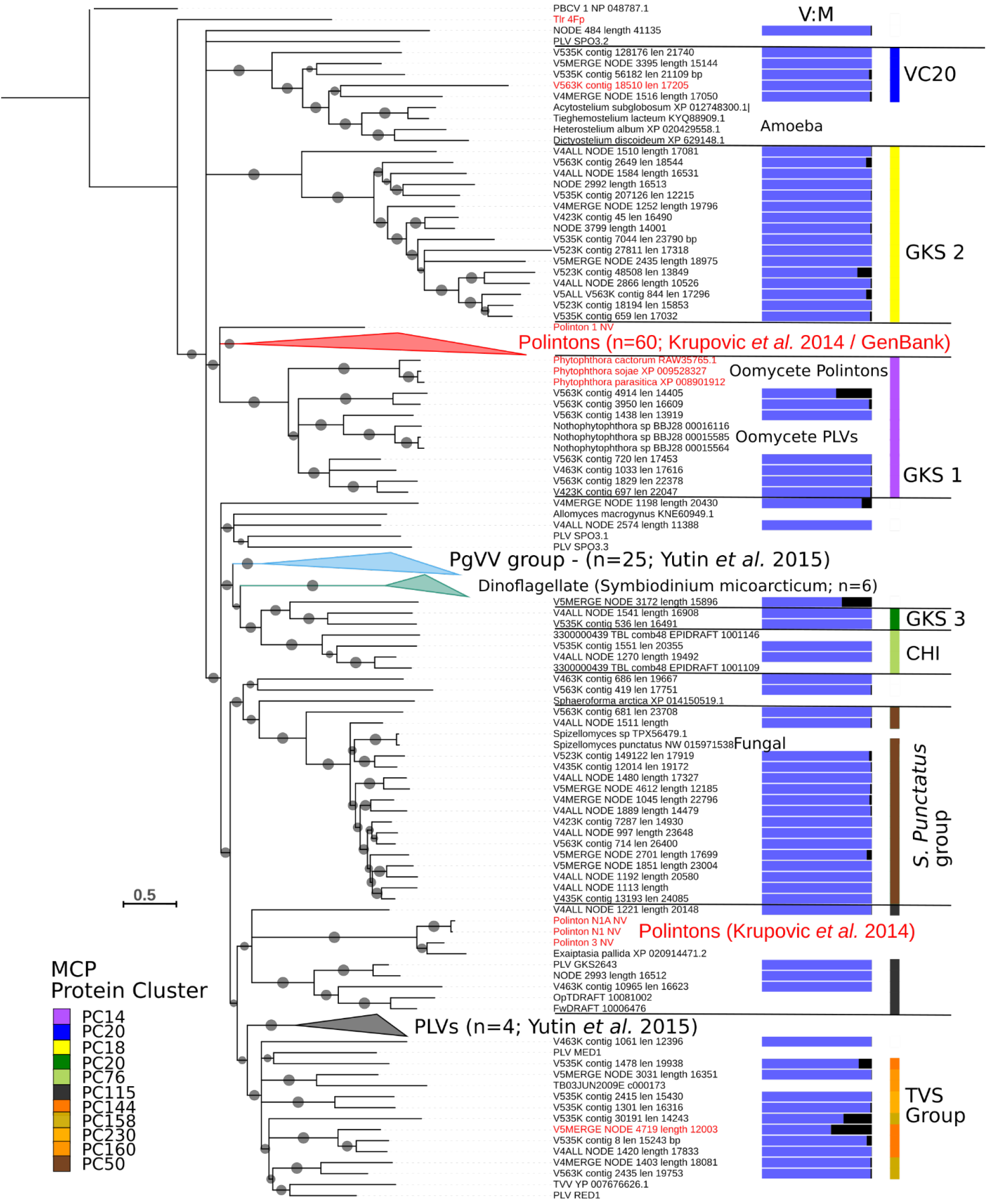
Phylogenetic analysis based on the Major Capsid Protein of Polinton-like viruses and Polintons. A maximum likelihood tree constructed using MCP genes from Gossenköllesee PLVs, and known Polintons. Grey circles represent SH (Shimodaira–Hasegawa)-like local support values of between 50-100%. Branches with <50% support were collapsed. Viruses with a confirmed Polinton genomic configuration (pPolB and RVE integrase) are labelled in red text. V:M represents the ratio of reads recruited from the viral (blue) and microbial (black) fraction metagenomes (Gossenköllesee only). The colour key represents the assigned protein cluster for the MCPs based on the network-based analysis. Annotated groups show the network-analysis based clusters from Figure 4, or previously defined MCP groups.

The majority of known Polinton MCP genes formed one large group, which also contained five MCP genes from Gossenköllese. However, from the PSI-BLAST searches we retrieved several new, related MCP genes from uncharacterized eukaryotic proteins in GenBank, which were also distinct from known Polintons. Six unique MCP genes were found in the dinoflagellate *Symbiodinium microadriaticum*, HHpred analysis of aligned MCP genes confirmed they were distantly related the MCP gene of PBCV-1 (96.6% probability) consistent with them being Polinton/PLV capsids. We also identified novel MCP members in the fungus *Allomyces macrogynus* and the Ichthyosporean *Sphaeroforma arctica*. In terms of known groups of PLVs, the *Tetraselmis striata* virus group (TVS group)^3^ was extended with 12 new MCPs from Gossenköllesee, whilst no new MCPs from our metagenomes fell into the *Phaeocyctis globosa*virus virophage (PgVV) group which contained algal associated PLVs, including the *Chrysochromulina parva* virus virophages.

PLVs were present in all three sampling times in Gossenköllesee (October 2017, February and April 2018), however their highest relative abundance occurred in the ice free sampling period (October) when up to 4.9% of all metagenomic reads from the virome recruited (mapped back to) to PLVs (Table S1). Several of the PLVs in Gossenköllesee were highly represented in the viromes, comparable to the most abundant bacteriophages, including one 17.8 kb circular contig (Figure 2) (PLV_GKS2643), which was the 8^th^ most abundant virus contig in the entire virome (relative abundance based on normalised depth of recruited reads), sequenced to a depth of over 1000 times coverage per Gb. Given the relatively small size of this genome, this particular PLV is likely a dominant member of the virus population. Relative metagenomic recruitments for the 83 PLVs vs 23 virophages (contigs >10kb) were 32 Mb vs 2 Mb per Gb metagenome, hence PLVs were more diverse and more abundant than the virophages (Table S1). Each of the top three most abundant PLVs individually recruited more reads than the combined virophage population. Whilst the virophages are known to form virus particles, including the Mavirus virophage^17^ with its Polinton like genomic configuration, only *Tetraselmis striata* virus has been observed producing virions^12^. To address if the new PLVs we found in Gossenköllesee are genuine free virus particles or integrated elements in eukaryotic genomes, we compared the number of normalised metagenomic reads that mapped to our PLVs from the viral and microbial size fractions. We found that the majority (84%) of PLVs in our database recruited 95% or more reads from the viral size fraction (Table S1) (median 190-fold enrichment in the viral size fraction; range 1 – 2816 fold). This was similar to virophages (median 108-fold enrichment; range 1-1078) supporting the idea that the majority of PLVs are genuine virus particles. Six contigs with low representation in the virus size fraction (0-5 × coverage Gb^-1^), however, were enriched in the microbial fraction. Five of these possessed an MCP gene which fell into the main Polinton group of known integrated elements (Figure 1) and in three of these, we detected pPolB and RVE integrase genes, suggesting these were Polintons capable of genomic integration (Table S1). Several further PLVs also had a relatively strong signal in both the viral and microbial size fractions, which would be expected if they are capable of a duel lifestyle of free virus particles and integrated elements, such as Mavirus. These included the PLVs V5MERGE_NODE_4719_len12003, V535K_contig_30191_len_14243, V5MERGE_NODE_3172_length_15896 which also contained one or both of the Polinton core genes (pPolB and RVE).

**Figure 2.**
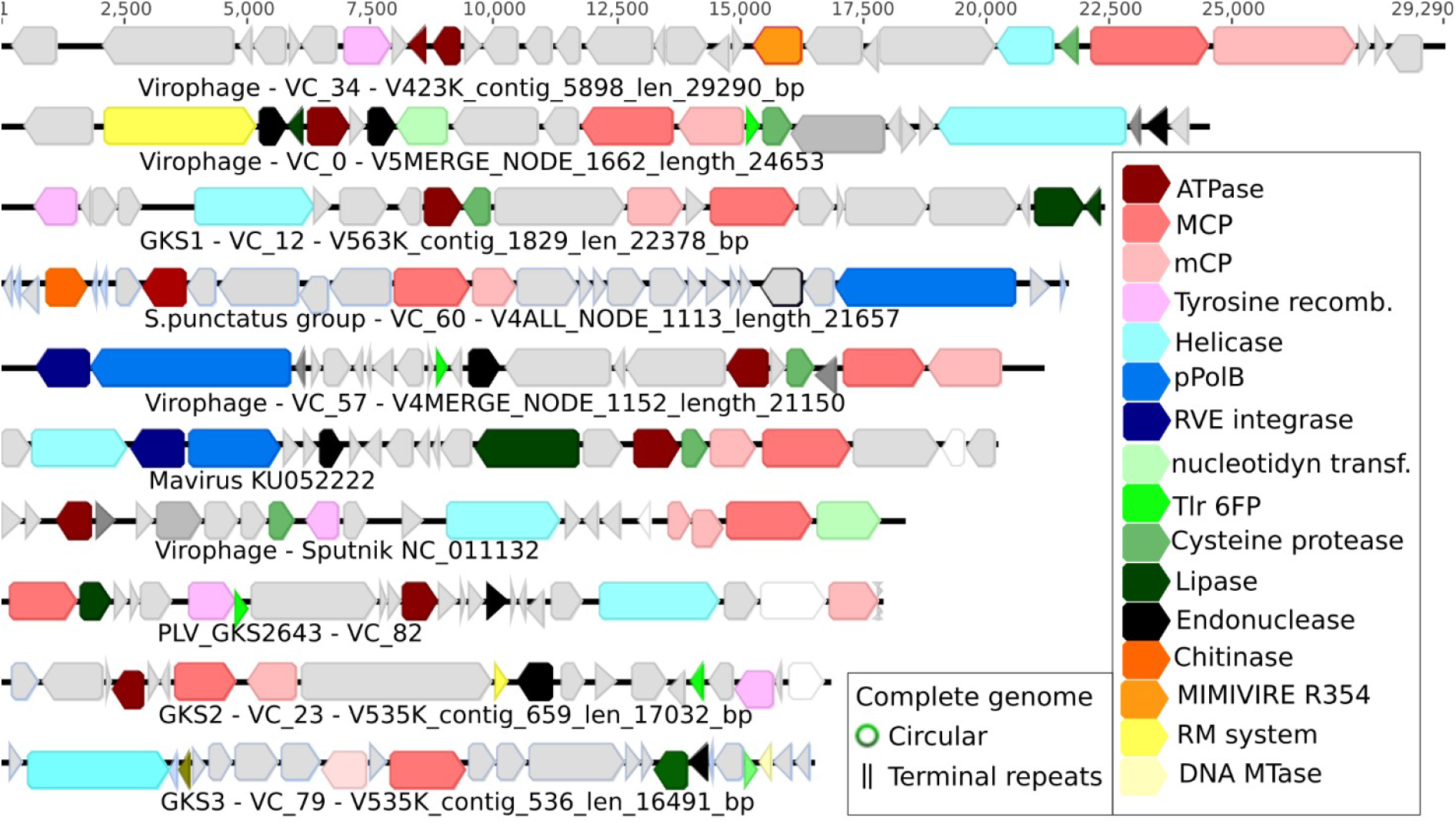
Genome maps of PLVs and virophages representing 7 of the major virus groups. The VC_34 virophage represents a virus cluster (n=12) of which all members possess a Cas4-like nuclease. The VC_57 virophage is a new Polinton/Mavirus type virophage with a virophage MCP and pPolB and RVE family integrase. PLV_GKS2643 was the most abundant PLV in Gossenköllesee. Approximately half of S. *punctatus* group members (VC_60) possessed chitinase gene.

Our viromes were generated from 0.2 μm filtered, DNase treated samples, which should remove most bacteria and digest contaminating (non-encapsidated) free DNA, however non-viral reads frequently remain in viromes despite extensive purification^18^. To assess the level of eukaryotic genomic contamination in the assembled virus-sized fraction, we searched all predicted genes against the SILVA SSU (16S/18S rRNA) database (BLASTP E-value < 10^-5^) to determine if contaminating eukaryotic DNA could account for all the observed PLVs. Only low-level eukaryotic contamination could be detected by this search across the viromes in 4 contigs (0.2-1 × coverage GB^-1^) (Table S4). The large relative abundance of PLVs in our dataset (up to 1198 × coverage Gb^-1^ and 4.9% of all reads) precludes the possibility that low-level eukaryotic DNA contamination could account for their detection. If the single most abundant PLV we found were integrated into a contaminating eukaryotic genome of 10 - 20 MB, the remainder of the genome would contaminate 30 - 60% of our total virome reads (assuming it is sequenced at a similar depth). Taking into account all the PLVs we discovered, the evidence strongly supports the conclusion they are virus particles. Namely, no PLVs were assembled into contigs longer than 42 KB (the maximum sized circular PLV we discovered); many could be circularized or exhibited terminal repeats; PLVs were abundant in the virus size fraction, hundreds of time more so than in the microbial fraction, and PLV were relatively more abundant than the virion-forming virophages.

To determine if the novel PLVs were part of a widespread larger group of viruses, we used all MCP genes as bait to interrogate publicly available metagenomes. Two profile Hidden Markov Models (profile HMMs) were built from the alignment of all new and known PLV and virophage MCP genes using HMMER (hmmer.org). These were used to interrogate the Integrated Microbial Genomes Virus (IMG/VR) database of globally derived viral sequences^19^ (retrieving 200 virophages and 167 PLVs) from terrestrial, marine and predominantly freshwater ecosystems (Figure 2). Selected publicly available metaviromes^20^ with numerous IMG/VR hits were further assembled (SPAdes) and interrogated (BLASTP; E-value cut off 10^-5^) retrieving 103 virophages and 376 PLVs. Hence this study detected or retrieved 335 new virophage and 625 new PLV genomes over 10kb. To complete the dataset, we added in 39 known virophages, 25 known PLVs at the time of analysis (Feb 2019) along with the 15 integrated elements from oomycetes and one from S. *punctatus*. Hence, we built a dataset of 1040 globally distributed viruses over 10kb in length, representing a 10-20-fold increase in known virophage and PLV genomes respectively.

To understand the relationships between the large numbers of virophages and PLVs in our dataset, we used a network-based analysis of shared protein clusters using vConTACT v.2.0^21^. Such an approach is suitable when faced with numerous distantly related viruses which may undergo frequent genetic exchange. This is because all genes are compared, uncovering new relationships and shared gene content which complements the MCP phylogeny. Network analysis grouped 739 viruses into 61 Virus Clusters (VCs) with ≥ 3 members. (in bacteriophages this is equivalent to genus level groupings, however this remains untested in eukaryotic viruses) and 64 viruses into VCs with only 2 members per cluster (Figure 4). A further 213 viruses were outliers and 24 were singletons (sharing no protein clusters and not included in the network), including *Tetraselmis striata* virus (TsV-N1)^12^ which infects a microalgae of the Chlamydomonadaceae family. This lack of clustering of singletons and outliers suggests a large-scale under-sampling of the true PLV diversity, as TsV-N1 MCP is part of the PLV group^3^ (Figure 1).

**Figure 3.**
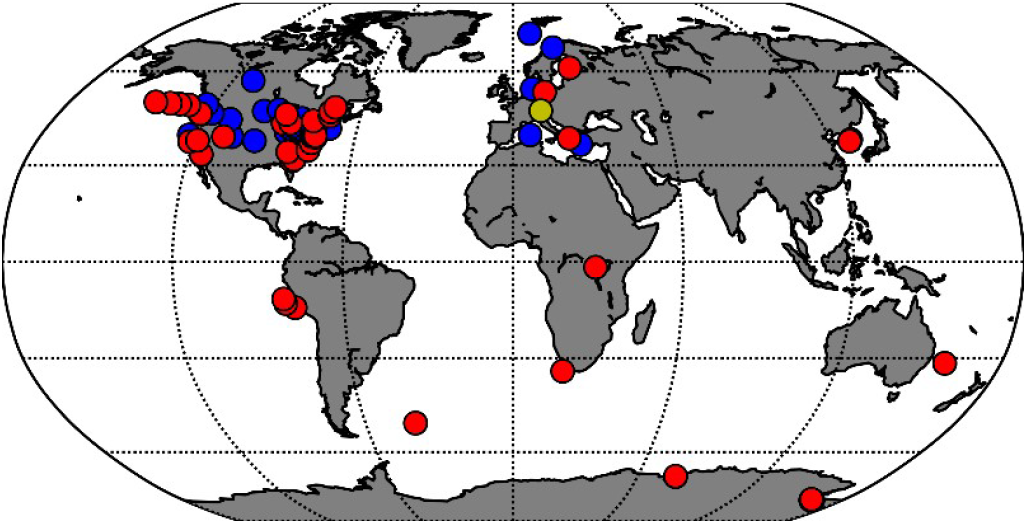
PLVs and virophages from widespread locations. Each point represents a location where one or more virus contigs were retrieved from the IMG/VR database, public metagenomes and Gossenköllesee. Colours represent: red - PLVs and virophages; blue-virophages only, yellow - Gossenköllesee PLVs and virophage.

**Figure 4.**
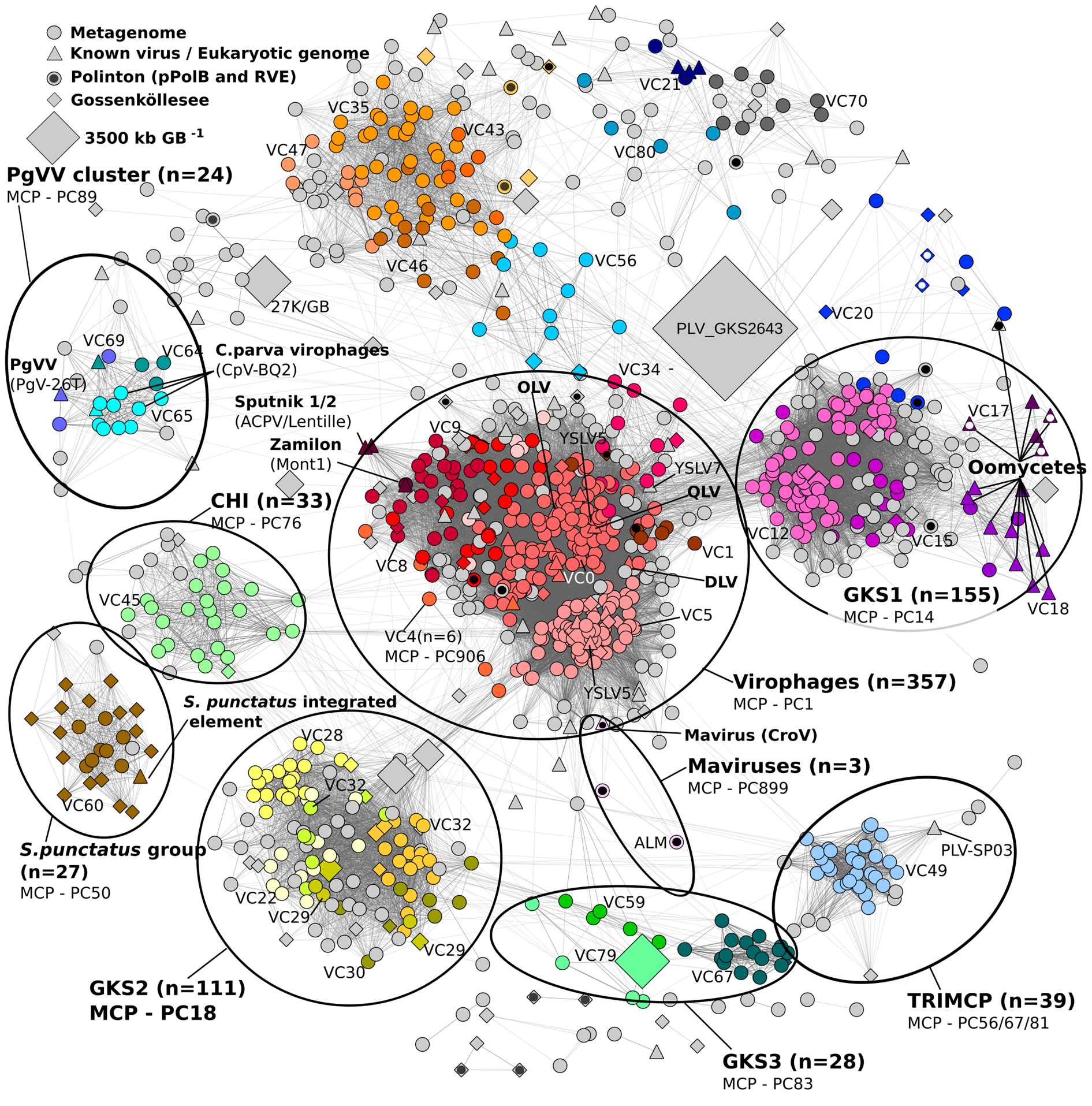
Polinton-Like Viruses (PLV), Polintons and virophages share a network of genes. A network-based analysis of shared protein clusters from 1015 virophage, PLV and Polinton genomes visualised in Cytoscape. Each symbol (node) represents a complete or near complete virus genomes of 10-42kb, the lines (edges) represent the strength of the connectivity (edge weight) between each genome. Symbols represent virus origin: Gossenköllesee metagenomes (diamond); isolated/previously detected viruses (triangle); retrieved from global metagenomes (circle). Polintons containing pPolB and RVE integrase include a black or white circle inside the symbol. The size of the diamond symbols represents the mean relative abundance of the virus in Gossenköllesee metagenomes. Virus clusters (VCs) with 5 or more members or with a known isolate are represented by colours and labelled. Known isolated viruses and integrated elements are labeled in bold with associated giant virus (if known) in brackets. Circles show larger virus groups sharing the same major capsid protein cluster. Viruses with no shared protein clusters (n=25) are not present in the network.

From the network (Figure 4), it was apparent that individual VCs and outliers formed larger scale virus groupings which were consistent with the MCP protein cluster they possessed (Table S3) and the MCP groups formed from the phylogenetic analysis (Figure 1). All viruses possessing a virophage type MCP (PC_0001) formed a single large group, composed of 11 distinct VCs with 267 members and 90 outliers (Table S3). Within this group, isolated virophages Sputnik^6^, Sputnik 2^22^ and Zamilon^23^, associated with the amoeba *Acanthamoeba polyphaga* clustered together (VC5, n=4), separating them from the largest group of virophages (VC0, n=111), which contained the metagenomically detected Organic Lake Virophage^24^ (OLV) and the Yellowstone Lake Virophages^25^ (YSLV) that putatively infect photosynthetic protists (Figure 4). Mavirus fell into a diverged cluster (VC3, n=3), having an MCP protein distinct from the other virophages^26^. Many new viruses contained genes which may be involved in countering host defence mechanisms. The majority of viruses in our dataset (n=632) encoded a DNA methyltransferase gene as previously described for PLVs and virophages, however, at least 7 virophages from VC_0 and VC_4 possessed putative restriction modification systems, previously hypothesized to aid giant viruses in their defense against virophages^5^. Similarly, 61 viruses in our dataset, across all virus groups (apart from the *S. Punctatus* and CHI groups) contained a predicted E3 ubiquitin-protein ligase. E3 ligases are known to counter host immune defenses in vertebrate viruses^27^, they have previously been detected in one other PLV^11^. Interestingly, there was a further group of 12 virophages from VC_34 which encoded a Cas4-like nuclease (Figure 2). Such a gene has previously been detected in Mimivirus and has been implicated in the proposed MIMIVIRE defense system^28^.

Despite their highly diverged capsid genes, there was considerable genetic overlap between the virophages and a large cluster of PLVs in the GKS-1 group (n=155; Figure 4). All GKS-1 members shared the same MCP protein cluster (PC_0014), however, two virus clusters within this group (VC12 and VC15) shared a similar tyrosine recombinase (PC_0000) with the virophages (Table S5). Most GKS-1 group members also possessed a cysteine protease (Figure 2), which is involved in virophage capsid maturation and has been notably absent from the PLV until now^3^. In addition, GKS-1 and virophage viruses also shared lipase (PC_0008) endonucleases (PC_0013) and phospholipase (PC_0010) protein clusters (Table S5). Of the 15 oomycete elements in GKS-1, four were Polintons, possessing an RVE type integrase and pPolB, whilst three more possessed one of the two genes, yet their MCP genes were distinct from known Polinton MCPs (BLASTP E-value > 0.1). Five further oomycete integrated elements possessed no Polinton marker genes, instead encoding a DNA helicase-primase suggesting they were PLV. In *Phytophthora cactorum* strain 10300 and *P. infestans* (T30-4), both PLV and Polinton forms were present in the same strain, suggesting opportunities for gene transfer between the two virus forms are frequently encountered.

The *S. punctatus* PLV clustered into the eponymous group (n=27) (Figure 4). This early diverging chytrid fungus is noted for the possession of motile zoospores^29^. Concurrently, 52% of the viruses in this group contained a chitinase gene (Table S5; Figure 2), which would be a necessary adaptation to penetrate a fungal cell wall, suggesting that members of this cluster are able in integrate into fungal genomes. Additionally, 78% of viruses from the CHI group (n=32) also contained a chitinase gene (Table S5). In terms of the known PLVs, network analysis confirmed the PgVV group which included PgVV and the *C. parva* virus virophages^11^ (Table 1), all of which share the same MCP protein cluster (PC_0089) and are associated with giant viruses infecting haptophytes (Table 1). No Gossenköllesee viruses fell into the PgVV cluster, however several viruses from global metagenomes were included. The TRIMCP group had one Gossenköllesee member and contained a unique PLV from a previous study which had three distinct MCP genes (PLV-SP03)^3^. All 39 members of the TRIMCP group possessed the same unusual configuration.

An outstanding question posed by these new PLVs is whether they are capable of independently infecting and replicating in a host, as described for *Tetraselmis striata* virus, or whether they associate with giant viruses. PgVV and the *C. parva* virus virophages were found either attached to, or inside giant virus virions^10,11^, suggesting they may co-infect eukaryotic hosts with giant viruses in a manner similar to some known virophages. These confirmed giant virus associated PLVs coupled with genetic overlap between many PLV groups to virophages (Figure 4) raises the possibility that many more of the PLVs identified here could be giant virus-associated entities. Whilst we detected nucleocytoplasmic large DNA viruses (NCLDV) MCP genes in our viromes, we did not detect any shared genes between these contigs and the PLVs, as was the case for organic lake virophage and its giant virus host^24^. Not did we detect broken paired end reads mapping to both giant virus and PLVs, as previously suggested to signify NCLDV genomic integration hotspots^10^. Hence until these PLVs are isolated with their eukaryotic hosts, this remains an open question. The bottleneck in eukaryotic host identification for this dataset is currently the lack of protist genomes available to analyse. This will be made easier with the advent of large scale protist sequencing projects, such as the Darwin Tree of Life project, where numerous protest genomes will become available to mine for integrated PLVs.

It has been speculated that PLVs may resemble the first eukaryotic viruses, representing a link between bacteriophages of the Tectiviridae family and the Polintons^3^, which in turn are thought to have diverged into Maviruses and the virophages^1^. With over a 10-fold increase in genomes provided by our study, it becomes more apparent that Polintons, PLVs and virophages share many overlapping genes and characteristics, with the distinction between the groups now largely based on MCP type (Figure 4) rather than difference in core gene content. For example, serval PLV groups shared more gene content with the virophages than they did with each other (Figure 4). We also detected 28 viruses in our network which fulfilled the definition of Polintons (pPolB, RVE, MCP, mCP and ATPase). These viruses populated at least 8 VCs, spread across the virophages, GKS1 cluster (including integrated Polintons in oomycete genomes), the TVS group and several outlier viruses (Figures 2 and 4). Several of these viruses are also present in both the microbial and viral size fractions (Table S1), suggesting possible host genomic integration similar to Mavirus/Polintons. Given that these distantly related virus groups contain members with both Polinton and PLV genomic configurations (Figure 1); that Polintons and PLVs integrate into the same hosts; and that protein cluster networks suggest multiple genes are shared between all groups (Figure 4), it seems likely that frequent gene exchange created the two virus configurations across the groups. Hence, Polintons in microbial eukaryotes, virophages and PLVs appear to be diverged variants of a wider group, which represents an expansive assemblage of viruses, widespread in aquatic ecosystems.

This study establishes PLVs as an important component of many freshwater viromes, which given their diversity, likely infect an even wider range of hosts than we have currently identified. The majority of new PLV genomes in our dataset are largely based on an extensive analysis of a single alpine lake virome, using the new MCP genes to interrogate global metagenomes. It is therefore likely that as new viruses are discovered in different environments and MCP genes are identified, many more genomes will be uncovered from uncharacterised proteins in sequencing datasets. Hence, the PLVs we present here probably represent only a small fraction of their true global diversity.

---

**Figure S1.**
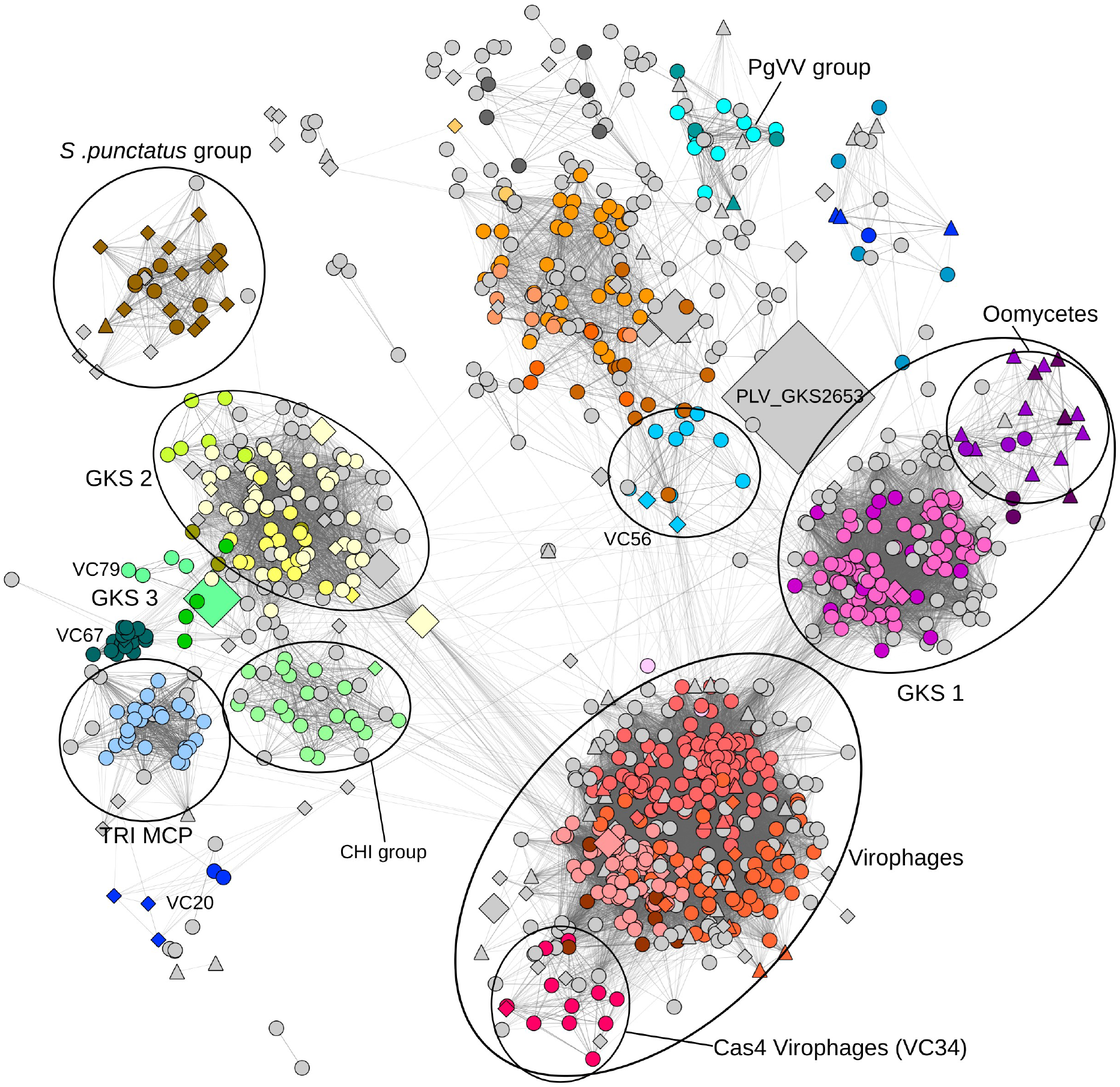
vConTACT2 network generated using BLASTP (E-value cutoff 10^-7^) proteinprotein comparisons. See Figure 4 for full description.

**Figure S2.**
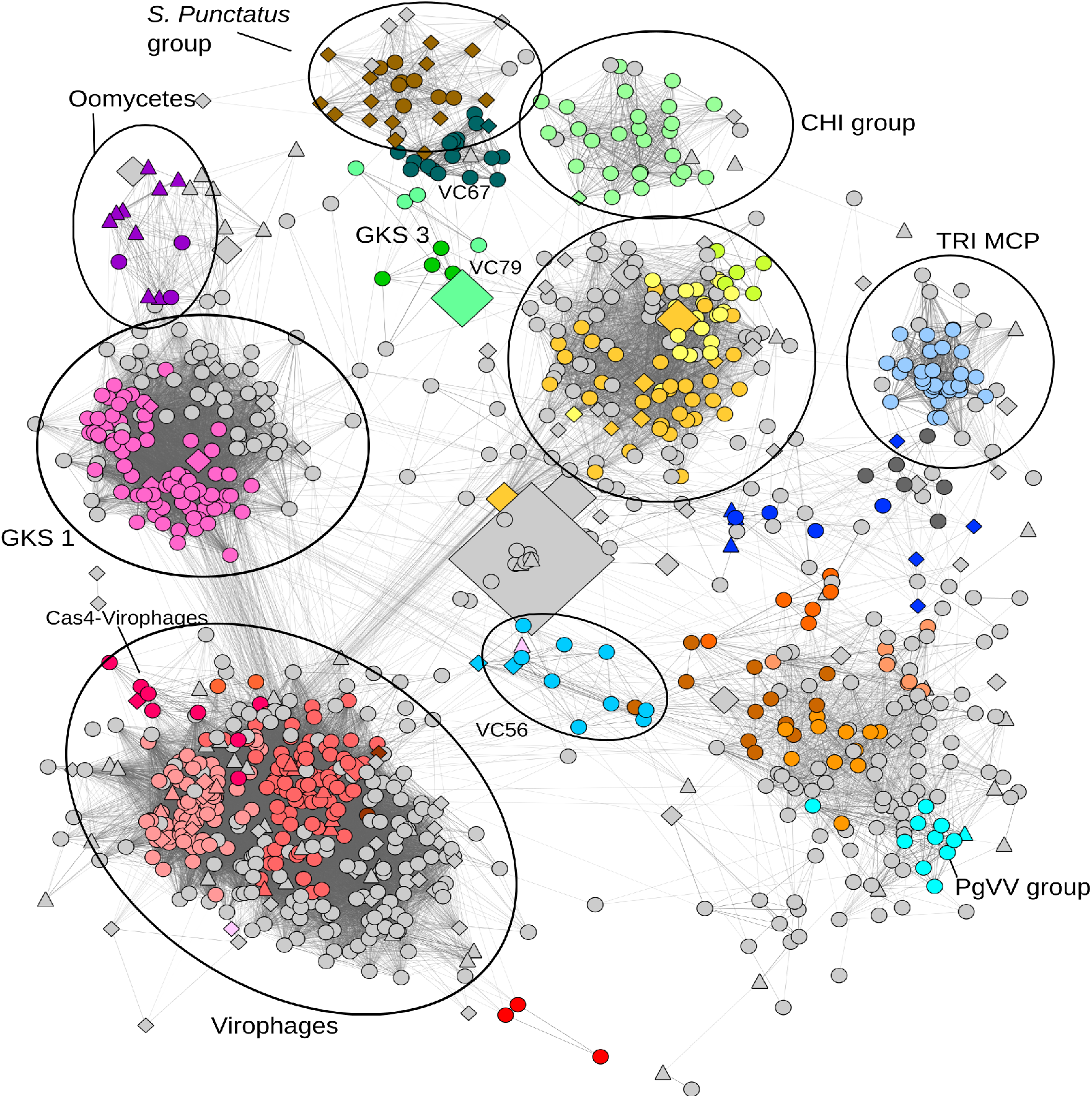
Higher stringency vConTACT2 network generated using BLASTP (E-value cutoff 10^-9^) protein-protein comparisons. See Figure 4 for full description.

## Methods

### Sampling and metagenome generation

We sampled an oligotrophic, high mountain lake (2417 m), Gossenköllesee (GKS, 47°13’46.7”N 11°00’47.9”E) in October 2017, February and April 2018 to generate paired virus and microbial size fraction metagenomes. For the virus metagenomes and for each sampling time, 40 L composite lake water samples (n=3) were collected from 1-8 m depth and passed in sequence through a 142mm GF/A (Whatman) then 142 mm 0.22 μm PES filter (Millipore). Virus concentration and purification was by iron chloride precipitation^30^, using pH 6 ascorbic acid to digest the iron-virus precipitate, 4 × Promega Wizard® Minicolumns per replicate with a ratio of 0.5:1 virus concentrate to Wizard® PCR Preps DNA Purification Resin (Promega: A7170). The resultant extract was found to be inhibitory to all enzymatic reactions, including Illumina library preparations, so it was further cleaned prior to sequencing. The crude extract in TE buffer (~800 ml per replicate) was divided into two, ethanol precipitated using 0.3 M NaCl and 2μl Glycogen (Invitrogen 10814010) and resuspended in 100 μl 1 × TE buffer (pH 8). The concentrate was then washed via diafiltration using Amicon® Ultra-2 mL centrifugal filters with a 30 kilodalton cutoff (Merk UFC203024). Diafiltration consisted of adding 2 ml of wash buffer and spinning down at 2000 × g until the sample volume reached ~100 μl. The filtrate was discarded and the wash repeated. The diafiltration steps were 3 × rounds of wash buffer 1 (100mM EDTA, 300mM NaCl, 10 mM Tris-Cl pH 8); 2 × rounds of wash buffer 2 (300 mM NaCl and 10 mM Tris-Cl pH 8) and 2 rounds of 1 ×TE buffer (pH 8). The resultant washed DNA (100 μl) was then cleaned with a DNeasy PowerClean Pro Cleanup Kit (Qiagen 12997-50).

To extract DNA from the microbial size fraction, 1 litre of composite lake water sample per timepoint (n=1) was filtered onto a 47 mm, 0.2 μm Polycarbonate membrane (Whatman) which was frozen until further processing. DNA was extracted from the filter (Acinas lab DNA extraction method)^31^ and purified by a standard Phenol:Chloroform:Isoamyl Alcohol (25:24:1, v/v) extraction with the modification of 3 × Chloroform:Isoamyl alcohol washes (24:1). Residual chloroform was removed from the sample by cleaning with a DNeasy PowerClean Pro Cleanup Kit (Qiagen).

Sequencing libraries for all samples were prepared by the Bristol Genomics Facility (http://www.bristol.ac.uk/biology/genomics-facility/) using the TrueSeq nano LT library prep kit and sequenced on an Illumina NextSeq 500 using 2 × 150bp paired end reads on two high output runs.

### Assembly and gene prediction

Raw reads were trimmed for Illumina adapters using Trimmomatic^32^ before each replicate metagenome was assembled separately using SPAdes^33^ version: 3.12.0 (settings: --meta-k 21,33,53,77). A pooled assembly for each virome was also created for each time-point by concatenating the 3 replicate fasta files into one assembly. Genes were predicted on all assembled virus contigs >5kb using GeneMarkS^34^ Putative complete genomes, i.e. circular contigs, or those with terminal direct or inverted repeats were identified via read mapping with Bowtie2^35^ (--sensitive --no-unal -I 0 -X 800). The resulting SAM file was searched for contigs where discordantly mapping reads over 10 kb were within 1000bp of each end of the contig AND their coverage depth was >10% of the mean contig coverage. We flagged such contigs as a circular genome or a genome possessing terminal repeats.

### Detecting virophages and PLVs in our metagenomes

To detect virophages in our dataset, we downloaded all virophage MCP genes available from GenBank (Dec 2018) and concatenated these with those from two other recent metagenomic studies^26,36^. This file was used to interrogate the predicted genes from the pooled metagenomic assembles (BLASTP E-value cutoff 10^-5^). Amino acid sequences from a subsample of these putative virophages (n=13; Table S6) were annotated by HHpred^14^ for remote protein homology detection (https://toolkit.tuebingen.mpg.de/#/tools/hhpred). Genes from these confirmed virophages were then pooled with all known virophage genes to create a custom database which was used to search all our metagenomic assemblies for further distantly related virophage-like contigs (BLASTP E-value cutoff 10^-5^). A virophage-like virus was flagged where the following criteria were all met: 1) ≥20% of predicted genes on a metagenomic contig hit to the custom virophage gene database; 2) The contig was between 10 and 45kb in length; 3) No virophage MCP was detected. The most abundant, complete (circular) virophage-like genome was then selected to be annotated by HHpred searches against the Protein Data Bank, where we identified an ATPase, minor Capsid Protein (mCP) and an MCP gene with distant homology to that of the giant virus *Paramecium bursaria Chlorella* virus 1 (PBCV-1), as has been reported for Polintons and Polinton-like viruses^16^. This confirmed MCP gene was then used to re-interrogate our virophage-like virus dataset using BLASTP to identify further similar capsid genes (E-value cutoff 10^-5^). We then repeated this procedure on the remaining virophage-like virus contigs (Table S2), identifying a double jelly-roll fold capsid related to PBCV-1 or Faustovirus, and retrieving related genomes until we had either found a capsid or annotated via HHpred all virophage-like virus contigs. Finally, to further improve our detection of PLV in our metagenomes we also gathered all contigs that contained an MCP gene homologous to any known Polinton or PLV from the literature (BLASTP E-value <10^-5^). In some cases, the MCP gene could still not be detected in a putative PLV after multiple searches, however the presence of an ATPase and an mCP allowed us to carry these putative PLV contigs forward in the analysis and retrieve related contigs from our metagenomes using the mCP gene as above. Once we had clustered the viruses (see below), the MCP genes were then identified via a more sensitive HHpred query using an alignment of a core protein. Contigs were confirmed as virophage or PLV when they met the following criteria: 1) 20% or more genes hit to a known or annotated virophage/PLV from Table S2 or Table S5 (BLASTP E-value <10^-5^); 2) contigs were between 10-42kb; 3) an MCP gene was found. These settings were the result of extensive testing and annotation checks by HHpred to ensure no false positives were produced. As we retrieved viruses from multiple metagenomes, the same viruses was often retrieved multiple times, redundancy was therefore removed from the dataset using CD-HIT^37^ (Settings: psi-cd-hit.py - c 0.7 -G 1-circle 1-prog megablast) to produce a non-redundant PLV fasta file. The same procedure was repeated to remove redundancy from the matching virophage contigs.

### Public metagenome searches

To determine if the Gossenköllesee PLVs were unique to high mountain lakes, or part of a globally dispersed group of viruses, we interrogated large metagenomic datasets using the novel MCP genes as bait. The complete IMG/VR protein dataset^38^ was downloaded (IMG_VR_2018-07-01_4; Dec 2018) along with all metadata (https://img.jgi.doe.gov/vr/). All known PLV MCP genes from GenBank and the literature (n=25; Table 1) were pooled with all identified PLV MCPs from Gossenköllesee (n=83) and aligned using MUSCLE^39^. A profile Hidden Markov Model (profile HMM) was built from the alignment using the hmmbuild command of HMMER (hmmer.org). This profile HMM was used to interrogate the downloaded IMG/VR protein dataset for PLV MCPs (hmmsearch --noali -E 0.00001). All matching records were retrieved from the IMG/VR fasta file (IMGVR_all_nucleatides.fna) as full contigs. For the virophage MCPs, an identical approach was used to create a separate profile HMM and the matching contigs were pooled with the PLV fasta file. Many IMG/VR PLV scaffolds, particularity circular ones, were redundant. This was removed via two rounds of clustering with CD-HIT (initial round settings: psi-cd-hit.py -c 0.9 -G 0 -aL 0.2 - circle 1 -prog megablast; second round: psi-cd-hit.py -c 0.7 -G 1 -circle 1 -prog megablast). Many IMG/VR virophage and PLV hits were found in two large scale sequencing projects from freshwater viromes from South Korea, Han River (PRJEB19373) and Lake Soyang (PRJEB15535)^40^. To enhance hits for matching viruses from these viromes, we downloaded all trimmed metagenomic reads from these two BioProjects from the European Bioinformatics Institute (EMBL-EBI; https://www.ebi.ac.uk/), re-assembled each individual metagenome and cross assembled 3 metagenomes (PRJEB19373) using SPAdes (settings: --meta-k 21,33,53,77). Virophage and PLVs were retrieved, made non-redundant and confirmed as before. To interrogate global marine metagenomes for similar PLV, we downloaded all Tara Oceans assemblies^41^ (<0.22 μm) from EMBL-EBI (https://www.ebi.ac.uk/ena/about/tara-oceans-assemblies) and predicted genes as before. We used all PLV MCP genes as a BLAST query (BLASTP E-value < 10^-5^) against the Tara database as above. However, no new PLVs were identified, with only those identified in a previous study^3^, hence already in our dataset, being detected.

### Network analysis

To assess the relatedness of the PLVs and virophages, we used a network-based approach to cluster virus sequences based on shared protein clusters (PCs) on each contig. Using all 1040 PLVs and virophages, protein clustering and generation of virus clusters (VCs) was performed by vConTACT v.2.03^21^ (settings: BLASTP 1e-4) on the CyVerse platform (http://www.cyverse.org/). The network was visualized using Cytoscape version 3.7.0 using an edge weighted forced spring embedded layout. The BLASTP protein clustering threshold was chosen to allow grouping of all distantly related virus group, however, two additional network analyses were conducted using stricter BLASTP protein-protein clustering thresholds, E-value 10^-7^ (Figure S1) and 10^-9^ (Figure S2) to assess the validity of the results.

### Protein cluster annotation

Owing to the novelty of the virus genomes in our dataset, most virus genes displayed little or no homology to known genes using BLASTP searches against the GenBank non-redundant protein database. More sensitive searches were carried out using Hidden Markov Model (HMM) comparisons to annotate protein clusters. The output from *vConTACT v.2.0* grouped the amino acid sequences from 1040 viruses into 2213 protein clusters based on sequence homology. These were annotated as follows: 1) Members from each protein cluster were aligned using MUSCLE; 2) a profile HMM was built from each alignment using HMMER (hmmer.org) 3) the profile was used to interrogate UNICLUST30 (uniclust30_2018_08) and Swiss-Prot (https://www.uniprot.org/) databases (HMMSEARCH E-value cutoff 10^-4^). The top protein hit from each search was picked (excluding any hits to “Uncharaterized protein”) to create an annotation file (Table S7). For the most abundant protein clusters and the core genes from each virus cluster, MUSCLE alignments were used to search the Protein Data Bank (PDB_mmCIF70_10_Apr) using the HHpred server for remote protein homology.

### Phylogenetic analysis

MCP genes from Gossenköllesee were combined with known PLVs and Polintons from previous studies. To retrieve distantly related MCP homologues, we searched Gossenköllesee MCPs against the GenBank non-redundant protein database (BLASTP E-value <10^-5^). Where a GenBank hit was found we confirmed this as an MCP by HHpred annotation, then combined the hits into a PSI-BLAST iterative search. In this way we detected MCP homologs in distantly related eukaryotic genomes. Alignment of PLV MCPs was by MAFFT version 7 (E-INS-i iterative refinement method)^42^. Maximum likelihood analysis was performed using PhyML^43^ (LG substitution model with Shimodaira–Hasegawa-like estimation of branch support), the tree was visualised in iTOL v4 (https://itol.embl.de/) using PBCV-1 as an outgroup.

### Data Availability

The following data is available at https://figshare.com/s/c1557d3e5835d2928214. All nucleotide sequences from the PLV and virophages sequenced in this study from Gossenköllesee plus those reassembled from EBI metagenomes or retrieved from IMG/VR; all amino acid gene predictions used for virus clustering; vConTACT protein clusters; alignments of MCP genes. Profile HMMs for PLV and virophage MCPs.

All IMG/VR derived sequences listed in Table S3 are also available for download at https://img.jgi.doe.gov/vr/.

European Bioinformatics Institute metagenomes are available at https://www.ebi.ac.uk/metagenomics/ using the Bioproject IDs in Table 1.

## Acknowledgements

We thank Declan Schroeder for early advice on the manuscript, Birgit Sattler for assistance with sampling, Christopher Buck for stimulating discussion on animal Polinton-like viruses and the Bristol Genomics Facility for help with optimising DNA library preparation and sequencing. The computational results presented have been achieved (in part) using the HPC infrastructure LEO of the University of Innsbruck. This study was supported by a grant from the Austrian Science Fund (FWF, M 2299-B32).

## Author Contributions

C.B and R.S designed the study. C.B performed the sampling, metagenome generation and data analysis. C.B and R.S wrote the manuscript.

## References

1. Krupovic, M. & Koonin, E. V. Polintons: A hotbed of eukaryotic virus, transposon and plasmid evolution. Nat. Rev. Microbiol. 13, 105–115 (2015).

2. Koonin, E. V., Dolja, V. V. & Krupovic, M. Origins and evolution of viruses of eukaryotes: The ultimate modularity. Virology 479–480, 2–25 (2015).

3. Yutin, N., Shevchenko, S., Kapitonov, V., Krupovic, M. & Koonin, E. V. A novel group of diverse Polinton-like viruses discovered by metagenome analysis. BMC Biol. 13, 1–14 (2015).

4. Krupovic, M., Kuhn, J. H. & Fischer, M. G. A classification system for virophages and satellite viruses. Arch. Virol. 161, 233–247 (2016).

5. Duponchel, S. & Fischer, M. G. Viva lavidaviruses! Five features of virophages that parasitize giant DNA viruses. PLOS Pathog. 15, e1007592 (2019).

6. La Scola, B. et al. The virophage as a unique parasite of the giant mimivirus. Nature 455, 100–4 (2008).

7. Fischer, M. G. & Hackl, T. Host genome integration and giant virus-induced reactivation of the virophage mavirus. Nature 540, 288–291 (2016).

8. Fischer, M. G. & Suttle, C. a. A virophage at the origin of large DNA transposons. Science 332, 231–4 (2011).

9. Zhou, J. et al. Diversity of virophages in metagenomic data sets. J. Virol. 87, 4225–36 (2013).

10. Lescot, M. et al. Genome of Phaeocystis globosa virus PgV-16T highlights the common ancestry of the largest known DNA viruses infecting eukaryotes. Proc. Natl. Acad. Sci. 110, 10800–10805 (2013).

11. Stough, J. M. A. et al. Genome and Environmental Activity of a Chrysochromulina parva Virus and Its Virophages. Front. Microbiol. 10, 1–12 (2019).

12. Grébert, T., Pagarete, A., Sandaa, R.-A., Bratbak, G. & Stepanova, O. Tsv-N1: A Novel DNA Algal Virus that Infects Tetraselmis striata. Viruses 7, 3937–3953 (2015).

13. Hofer, J. S. & Sommaruga, R. Seasonal dynamics of viruses in an alpine lake: importance of filamentous forms. Aquat. Microb. Ecol. 26, 1–11 (2001).

14. Zimmermann, L. et al. A Completely Reimplemented MPI Bioinformatics Toolkit with a New HHpred Server at its Core. J. Mol. Biol. 430, 2237–2243 (2018).

15. Berman, H., Henrick, K. & Nakamura, H. Announcing the worldwide Protein Data Bank. Nat. Struct. Mol. Biol. 10, 980 (2003).

16. Krupovic, M., Bamford, D. H. & Koonin, E. V. Conservation of major and minor jelly-roll capsid proteins in Polinton (Maverick) transposons suggests that they are bona fide viruses. Biol. Direct 9, 1–7 (2014).

17. Born, D. et al. Capsid protein structure, self-assembly, and processing reveal morphogenesis of the marine virophage mavirus. Proc. Natl. Acad. Sci. U. S. A. 115, 7332–7337 (2018).

18. Roux, S., Krupovic, M., Debroas, D., Forterre, P. & Enault, F. Assessment of viral community functional potential from viral metagenomes may be hampered by contamination with cellular sequences. Open Biol. 3, 130160 (2013).

19. Chen, I. M. A. et al. IMG/M v.5.0: An integrated data management and comparative analysis system for microbial genomes and microbiomes. Nucleic Acids Res. 47, D666–D677 (2019).

20. Moon, K., Kang, I., Kim, S., Kim, S. J. & Cho, J. C. Genome characteristics and environmental distribution of the first phage that infects the LD28 clade, a freshwater methylotrophic bacterial group. Environ. Microbiol. 19, 4714–4727 (2017).

21. Bin Jang, H. et al. Taxonomic assignment of uncultivated prokaryotic virus genomes is enabled by gene-sharing networks. Nat. Biotechnol. (2019). doi:10.1038/s41587-019-0100-8

22. Desnues, C. et al. Provirophages and transpovirons as the diverse mobilome of giant viruses. Proc. Natl. Acad. Sci. 109, 18078–18083 (2012).

23. Gaia, M. et al. Zamilon, a novel virophage with Mimiviridae host specificity. PLoS One 9, 1–8 (2014).

24. Yau, S. et al. Virophage control of antarctic algal host-virus dynamics. Proc. Natl. Acad. Sci. U. S. A. 108, 6163–8 (2011).

25. Zhou, J. et al. Three novel virophage genomes discovered from yellowstone lake metagenomes. J. Virol. 89, 1278–85 (2015).

26. Roux, S. et al. Ecogenomics of virophages and their giant virus hosts assessed through time series metagenomics. Nat. Commun. 8, (2017).

27. Zhang, Y., Li, L. F., Munir, M. & Qiu, H. J. RING-domain E3 ligase-mediated host-virus interactions: Orchestrating immune responses by the host and antagonizing immune defense by viruses. Front. Immunol. 9, 1–12 (2018).

28. Levasseur, A. et al. MIMIVIRE is a defence system in mimivirus that confers resistance to virophage. Nature 531, 249 (2016).

29. Russ, C. et al. Genome Sequence of *Spizellomyces punctatus*. Genome Announc. 4, 4–5 (2016).

30. John, S. G. et al. A simple and efficient method for concentration of ocean viruses by chemical flocculation. Environ. Microbiol. Rep. 3, 195–202 (2011).

31. Adriana Alberti et al. Viral to metazoan marine plankton nucleotide sequences from the Tara Oceans expedition. Sci. Data 1–20 (2017). doi:10.1038/sdata.2017.93

32. Bolger, A. M., Lohse, M. & Usadel, B. Trimmomatic: A flexible trimmer for Illumina sequence data. Bioinformatics 30, 2114–2120 (2014).

33. Bankevich, A. et al. SPAdes: a new genome assembly algorithm and its applications to single-cell sequencing. J. Comput. Biol. 19, 455–477 (2012).

34. Besemer, J. & Borodovsky, M. Heuristic approach to deriving models for gene finding. Nucleic Acids Res. 27, 3911–3920 (1999).

35. Langmead, B. & Salzberg, S. L. Fast gapped-read alignment with Bowtie 2. Nat. Methods 9, 357–359 (2012).

36. Yutin, N., Kapitonov, V. V. & Koonin, E. V. A new family of hybrid virophages from an animal gut metagenome. Biol. Direct 10, 1–9 (2015).

37. Fu, L., Niu, B., Zhu, Z., Wu, S. & Li, W. CD-HIT: Accelerated for clustering the nextgeneration sequencing data. Bioinformatics 28, 3150–3152 (2012).

38. Paez-Espino, D. et al. IMG/VR: A database of cultured and uncultured DNA viruses and retroviruses. Nucleic Acids Res. 45, D457–D465 (2017).

39. Edgar, R. C. MUSCLE: multiple sequence alignment with high accuracy and high throughput. Nucleic Acids Res. 32, 1792–1797 (2004).

40. Moon, K., Kang, I., Kim, S., Kim, S. J. & Cho, J. C. Genomic and ecological study of two distinctive freshwater bacteriophages infecting a Comamonadaceae bacterium. Sci. Rep. 8, 1–9 (2018).

41. Roux, S. et al. Ecogenomics and potential biogeochemical impacts of globally abundant ocean viruses. Nature 537, 689 (2016).

42. Katoh, K., Rozewicki, J. & Yamada, K. D. MAFFT online service: multiple sequence alignment, interactive sequence choice and visualization. Brief. Bioinform. 20, 1160–1166 (2019).

43. Guindon, S. et al. New algorithms and methods to estimate maximum-likelihood phylogenies: Assessing the performance of PhyML 3.0. Syst. Biol. 59, 307–321 (2010)., 307–321 (2010).

